# A 3D-printed hand-powered centrifuge for molecular biology

**DOI:** 10.1101/519835

**Authors:** Gaurav Byagathvalli, Aaron F. Pomerantz, Soham Sinha, Janet Standeven, M. Saad Bhamla

**Author notes:** authors contributed equally. Please address correspondence to M.S.B.

## Abstract

The centrifuge is an essential tool for many aspects of research and medical diagnostics. However, conventional centrifuges are often inaccessible outside of conventional laboratory settings, such as remote field sites, require a constant external power source, and can be prohibitively costly in resource-limited settings and STEM-focused programs. Here we present the 3D-Fuge, a 3D-printed hand-powered centrifuge, as a novel alternative to standard benchtop centrifuges. Based on the design principles of a paper-based centrifuge, this 3D-printed instrument increases the volume capacity to 2 mL and can reach hand-powered centrifugation speeds up to 6,000 rpm. The 3D-Fuge devices presented here are capable of centrifugation of a wide variety of different solutions such as spinning down samples for biomarker applications and performing nucleotide extractions as part of a portable molecular lab setup. We introduce the design and proof-of-principle trials that demonstrate the utility of low-cost 3D printed centrifuges for use in remote and educational settings.

## INTRODUCTION

The centrifuge is an indispensable piece of equipment for laboratories, with general applications ranging from DNA isolation to clinical diagnostics. Yet, conventional centrifuges are often inaccessible outside of established lab settings (such as remote field sites), require a constant external power source, and can be prohibitively costly for STEM-focused programs. Progress has been made in the field of frugal science [2, 3, 6], a new approach towards making scientific tools more accessible and transportable, but there are many devices that remain to be developed or are currently in developmental stages. It is crucial to produce new low-cost devices to ensure increased access to scientific tools and expand scientific research without inhibition from costs and accessibility. Recently, 3D-printing technology has emerged as a revolutionary method for the rapid development and production of cost-effective scientific and diagnostic tools [4].

Here we present the 3D-Fuge, a 3D-printed device based on the principles of the paperfuge [2] as a low-cost human-powered alternative to standard benchtop centrifuges. The paperfuge is a recently developed ultralow-cost (20 cents), human-powered centrifuge that can be useful for applications including blood separation and disease diagnostics (such as anemia and malaria). Although the paperfuge is capable of centrifuging samples at speeds of up to 125,000 rpm and exert centrifugal forces of 30,000 Gs, it is limited by the sample volume it can hold (20 *μ*L per capillary tube). The 3D-Fuge in this study addresses the paper-based centrifuge limitation by expanding the liquid volume capacity (up to 2 mL) of samples, thereby enabling applications for workflows in molecular biology such as nucleotide extractions. It is capable of holding and spinning down four samples from the size of capillary tubes and PCR tubes to nucleotide extraction spin column tubes, allowing for the centrifugation of a wide variety of different solutions. The production of this device is fairly inexpensive, can be produced by anyone with access to a 3D printer, and can reach hand-powered centrifugation speeds up to 6,000 rpm (Fig 1A, B, K). It is highly portable and requires no continuous access to electricity, making it easy to transport and utilize for various applications. In order to demonstrate the capability of this device to perform routine experiments without access to conventional laboratory equipment, we carried out nucleotide extractions under both lab and remote field conditions (Fig 1, Case 1), with yields comparable to conventional benchtop centrifuges. We then performed downstream experiments from the 3D-Fuge extractions, including long-range PCR amplification and real-time nanopore DNA sequencing. We also integrate this device with a novel chromoprotein analysis application by producing bacterial pellets for quantification as part of a high-school STEM-program experiment (Fig 1, Case 2). Through these studies, we demonstrate the usage of the 3D-Fuge in different parts of the world (from a rainforest in Peru to a public high-school in USA) to validate its broad applicability. Due to its low-cost and ease of use, the 3D-Fuge can be valuable for a range of areas including field research, disease-screening in developing countries, and science education.

**FIG. 1.**
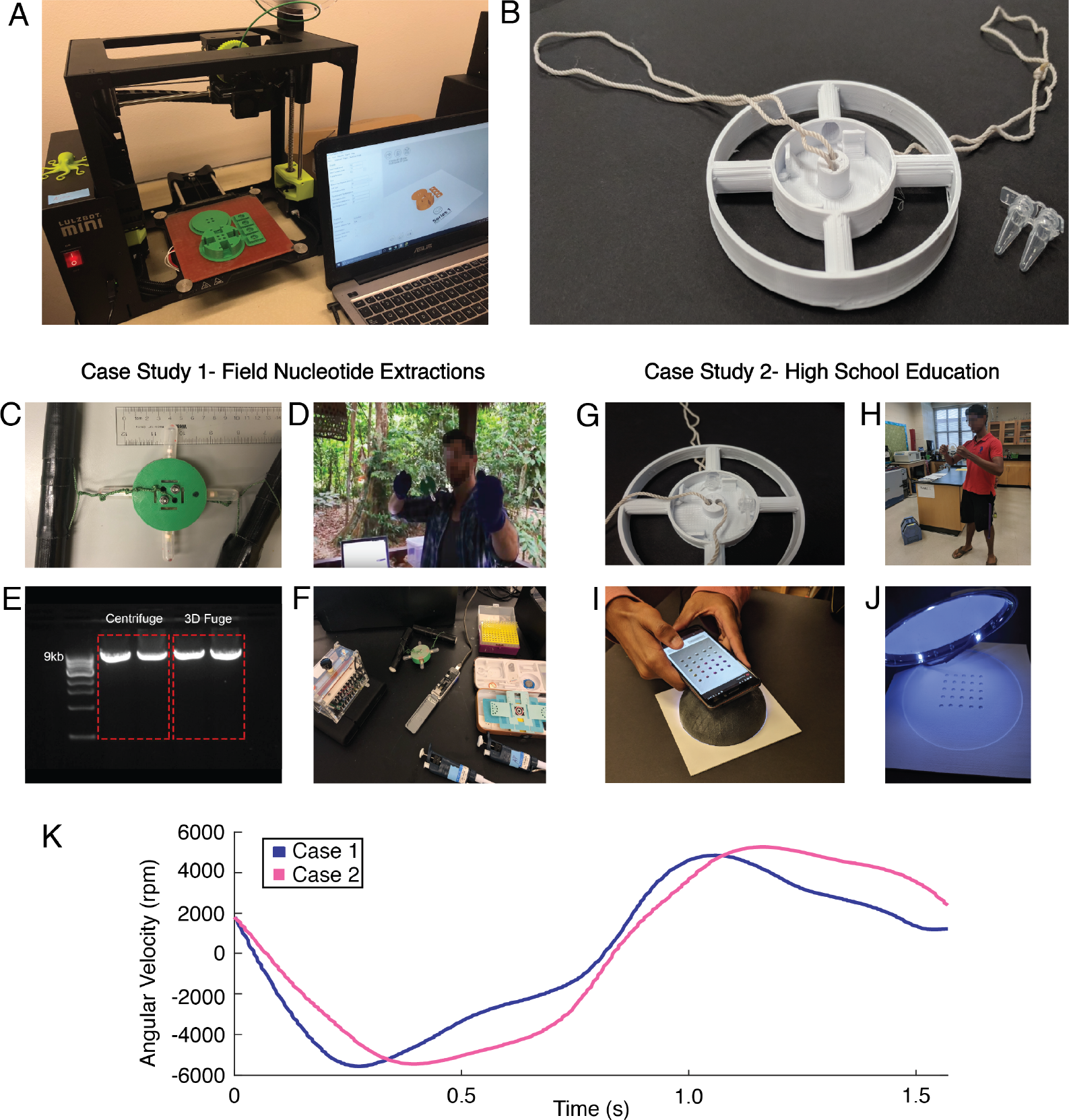
Overview of the 3D-Fuge. (A) Construction of the 3D-Fuge through 3D-Printing. (B) One of the 3D-Fuges utilized in this Case Study, with the 0.2 mL PCR Tubes placed alongside for comparison. (C) 3D-Fuge utilized in Case Study 1 (rainforest in Peru), shown holding up to four spin columns and flow-through tubes for nucleotide extractions. (D) Example usage of the 3D-Fuge in the field located in Southeastern Peru. (E) Comparison of long-range PCR results from human cheek swab nucleotide extractions using a standard benchtop centrifuge (left) and the 3D-Fuge (right). (F) Image of the portable lab setup, including the 3D-Fuge, the miniPCR (miniPCR) powered by an external battery pack, the MinION DNA sequencer (Oxford Nanopore Technologies), and the Foldscope (Foldscope Instruments). (G) Image of the 3D-Fuge utilized in Case Study 2 (High-School in Atlanta, Georgia) shown holding PCR tubes (200 *μ*L). (H) Usage of the 3D-Fuge in a high school. (I) Image of color output processing for data collection of chromoprotein expression. Phone captures the image of liquid culture pellets for RGB color analysis. (J) Sample illumination chamber utilized for standardized sample illumination. (K) Time evolution of the rpm of the 3D-Fuges over two cycles (counterclockwise and then clockwise inversion) utilized in each of the case studies. The peak rpm achieved is approximately 6,000 for both case studies. The above graphs illustrate the overall oscillatory motion of the 3D-Fuge, the change in angular velocity demonstrating the peak rpm over a given interval, and the changing relative centrifugal force with a relative maximum of 2100 g-force, through the given interval. **Authors Note:** Faces of individuals covered in accordance with **bioRxiv** policy to ‘avoid the inclusion of photographs and any other identifying information of people because verification of their consent is incompatible with the rapid and automated nature of preprint posting’.

## RESULTS

### Nucleotide Extraction and Nanopore Sequencing: Field trial with the 3D-Fuge in the Amazon Rainforest

Nucleotide extractions are a necessary first step for numerous molecular experiments, such as DNA sequencing projects, and often require centrifugation steps to separate and purify high-quality nucleic acids from the sample of interest. The ability to rapidly extract and purify nucleic acids with a low-cost hand-powered centrifuge can be useful for a wide range of molecular applications when one does not have access to conventional laboratory equipment, such as in the field or in resource limited settings. Portable sequencing projects are already emerging in applied field settings, including real-time species or environmental sample identifications [17, 18], pathogen diagnostics [8, 19]), and metagenomics [7, 12], but most studies thus far have utilized benchtop centrifuges with external power sources to prepare samples. Therefore, to demonstrate the capability of performing nucleotide extractions in a remote environment outside of a conventional laboratory, the 3D-Fuge was deployed during a biodiversity research expedition in Tambopata, Peru, at the Refugio Amazonas lodge (−12.874797, −69.409669, Fig 2, A, B).

**FIG. 2.**
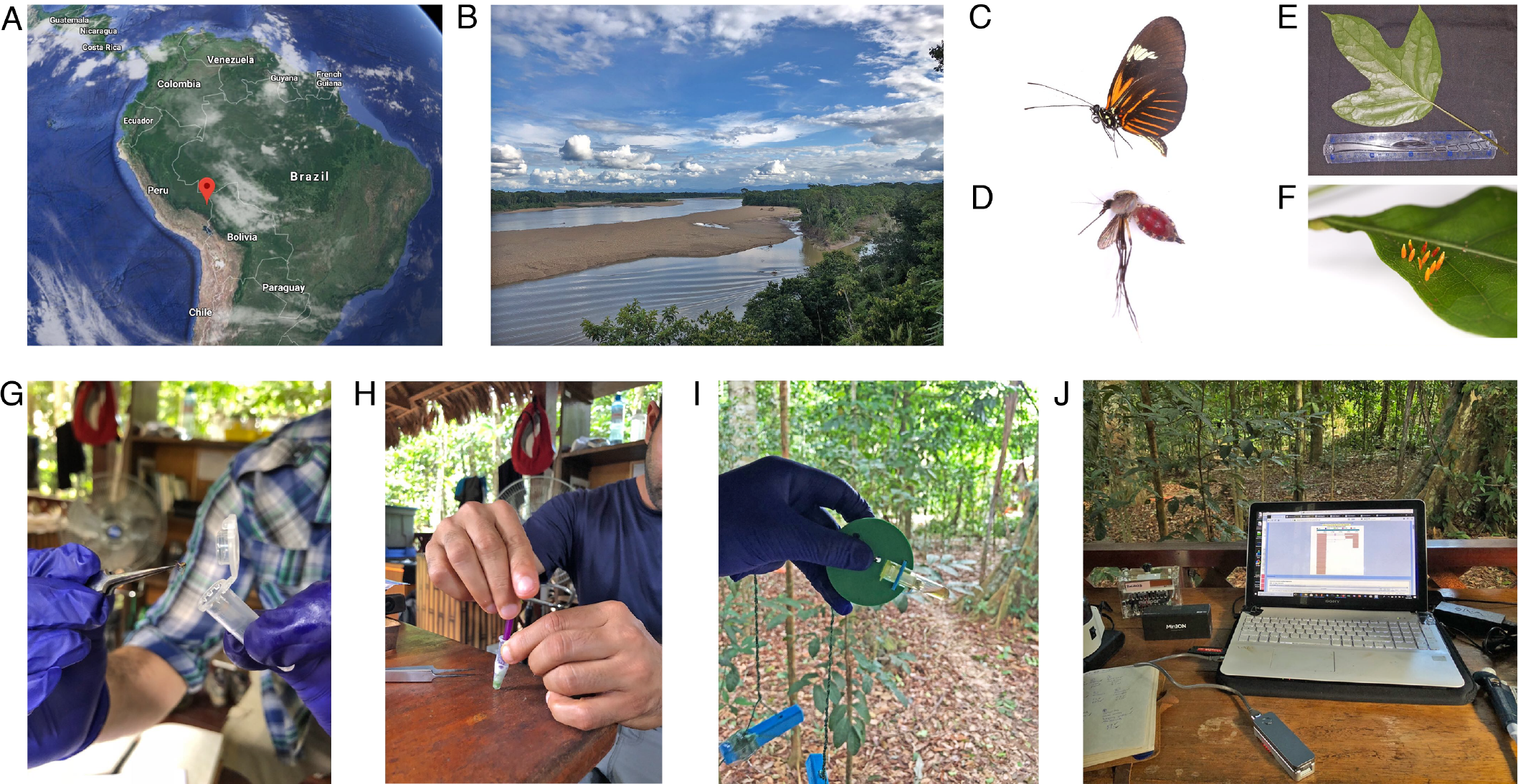
Overview of steps involved with sample collection, use of the 3D-Fuge for nucleotide extractions, and DNA sequencing in the Amazon rainforest. (A,B) Field site for the portable lab study located in Tamboapta, Peru. Examples of specimens collected for *in situ* DNA sequencing including (C) a butterfly, (D) a bloodfed mosquito, (E) a plant leaf, and (F) unknown insect eggs. (G,H) Tissue from specimens were homogenized and incubated in proteinase K and (I) the hand-powered 3D-Fuge was then used to perform DNA extraction and purification. (J) Purified DNA was subsequently used for downstream molecular steps including long-range PCR and real-time DNA sequencing on the MinION connected to a laptop. **Authors Note:** Face of individual covered in accordance with **bioRxiv** policy to ‘avoid the inclusion of photographs and any other identifying information of people because verification of their consent is incompatible with the rapid and automated nature of preprint posting’.

Before the expedition, we first compared DNA extractions in the lab using a standard benchtop centrifuge (Eppendorf, model 5415 D) and the hand-powered 3D-Fuge. A human cheek swab sample was collected and DNA extractions were carried out using the Quick-DNA Miniprep Plus Kit (Zymo Research, Irvine, CA, USA) according to manufacturer’s protocol. Eluted DNA yields were assessed using a Nanodrop and the results between centrifuge strategies, such as nucleotide concentration and A 260/280 ratio, were comparable (SI Fig 1B). The quality of the DNA from both extractions was also sufficient to perform long-range PCR amplification of 9,000 bp fragments of the human mitochondrial genome (Fig 1E), indicating that the hand-powered 3D-Fuge was capable of performing nucleotide extractions without requiring lab infrastructure.

Next, while in the Peruvian Amazon, specimens such as whole insects and plant leaves were collected and preserved in 1.5 *μ*L Eppendorf tubes containing DNA Shield (Zymo) for downstream processing (Fig 2 C-G). Molecular experiments in the field, including DNA extractions, long-range PCR, library preparation, and DNA sequencing, were performed using a highly miniaturized laboratory consisting of portable equipment [18]. The main components involved for this study include the 3D-Fuge, a small thermocylcer (miniPCR) and the MinION, a handheld nanopore-based sequencing device (Oxford Nanopore Technologies) (Fig 1F).

As before, all DNA extractions in the field were carried out using the Quick-DNA Miniprep Plus Kit (Zymo). Specimens were homogenized using a pestle and incubated with protinase K for 1-3 hours (Fig 2G, H). The 3D-Fuge for nucleotide extractions was designed specifically to fit and hold up to four standard 2 mL polypropylene spin-column tubes, which in this study were provided as part of the DNA isolation kit. Samples were transferred to the spin column tubes and placed into the 3D-Fuge which was spun by hand at maximum speed for approximately one to two minutes for each centrifugation step, including sample extraction, purification and elution (Fig 1D, Fig 2I). We were not able to quantify nucleotide extractions while in the field, so chose to use 1-3 *μ*L of eluted DNA from the 3D-Fuge for downstream long-range PCR reactions, which were performed using the Q5 Hot Start High-Fidelity 2X Master Mix (New England Biolabs, Ipswich, MA) and dual-indexed primers to amplify the ribosomal DNA (rDNA) cluster from plant and arthropod samples [13]. These PCR products were run on a gel to verify amplification after the expedition back at the lab and indicated all four extractions from the 3D-Fuge were successful (SI Fig 1C). In the field, approximately 1-2 *μ*L of PCR product for each of the samples were pooled and the Oxford Nanopore Technologies SQK-LSK 108 library preparation was carried out according to manufacturer’s protocol (ONT). The final library was run on the MinION and rDNA amplicons were sequenced in real-time on a laptop in the field (Fig 2J). Raw sequence reads were generated on a laptop in the field, and a bioinformatics pipeline was run to demultiplex samples and create a consensus sequence for each sample [13]. For the butterfly (Fig 2C), an rDNA consensus sequence of 4,658 bp in length was generated. The closest BLAST hit in the LepBase.org database was to *Heliconius*, a likely match based on morphology and genetic data. The bloodfed mosquito (Fig 2D) extraction yielded a consensus sequence 3,931 bp in length. BLAST and distance of tree results in the NCBI database yielded a closest match to a species in the genus *Psorophora* (SI Fig 1D). The plant specimen (Fig 2E) yielded a final consensus sequence 3,263 bp and the BLAST result was a closest match to species in the nightshade family Solanaceae, which was expected based on morphological identification by a botanist (Varun Swamy, personal communication). Finally, the eggs (Fig 2F) belonged to an unknown species of insect. The consensus sequence generated was 4,028 bp and BLAST results yielded a closest match to a butterfly species in the family Pieridae. Interestingly, the host plant was not detected in the sequence data with the insect eggs, but we did pick up a fungal sequence as well with a BLAST hit to the genus *Zymoseptoria*, which may have been an environmental sample on the leaf. Overall, the portable lab equipment enabled detection of specimens through rapid *in situ* DNA barcoding and use of the hand-powered 3D-Fuge demonstrated the feasibility of extracting high quality nucleic acids from samples in a remote tropical environment such as the Amazon rainforest.

### Chromoprotein Analysis for Synthetic Biology Education in High Schools

Reporter proteins are a quintessential part of synthetic biology [9], from identifying successful expression of genetic constructs to acting as biomarkers for diagnostic applications[11, 21]. Chromogenic proteins (or chromoproteins) are examples of such reporters capable of producing a color visible to the naked eye, unlike fluorescent proteins which are more commonly utilized [1, 15]. While fluorescent proteins are often quantified using plate readers, chromoproteins rely on the Red-Green-Blue (RGB) and Hue-Saturation-Luminance (HSL) color spaces for measurements [15], specifically with a focus on the corresponding hue value. In order to successfully obtain the hue values for bacterial samples expressing the reporter proteins, they must be concentrated into pellets to standardize the color values. Here, we report the usage of the 3D-Fuge for centrifugation of bacterial pellets for corresponding measurements using a sample illumination chamber and RGB color-analysis tool. This iteration is capable of centrifuging four 0.2 mL PCR tubes at speeds of up to 6,000 rpm, and to demonstrate its functionality, we centrifuge chromoprotein-expressing bacteria for color measurements.

The IPTG-inducible chromoprotein plasmid constructs (obtained from ATUM Bio, 2018)(Fig 3E) were transformed into DH5a *E. coli* and subsequently inoculated into liquid cultures with varying concentrations of Isopropyl *β*-D-1-thiogalactopyranoside (IPTG). Following the growth phase, liquid cultures were allowed to settle, and a portion of the settled particulate was centrifuged using the 3D-Fuge for 5 minutes. The supernatants were discarded and the obtained cellular pellets were were placed into the sample illumination chamber to capture an image with a phone. The image is then processed to extract the RGB/HSL values to determine the hue of the bacterial pellet (Fig 3B, D).

**FIG. 3.**
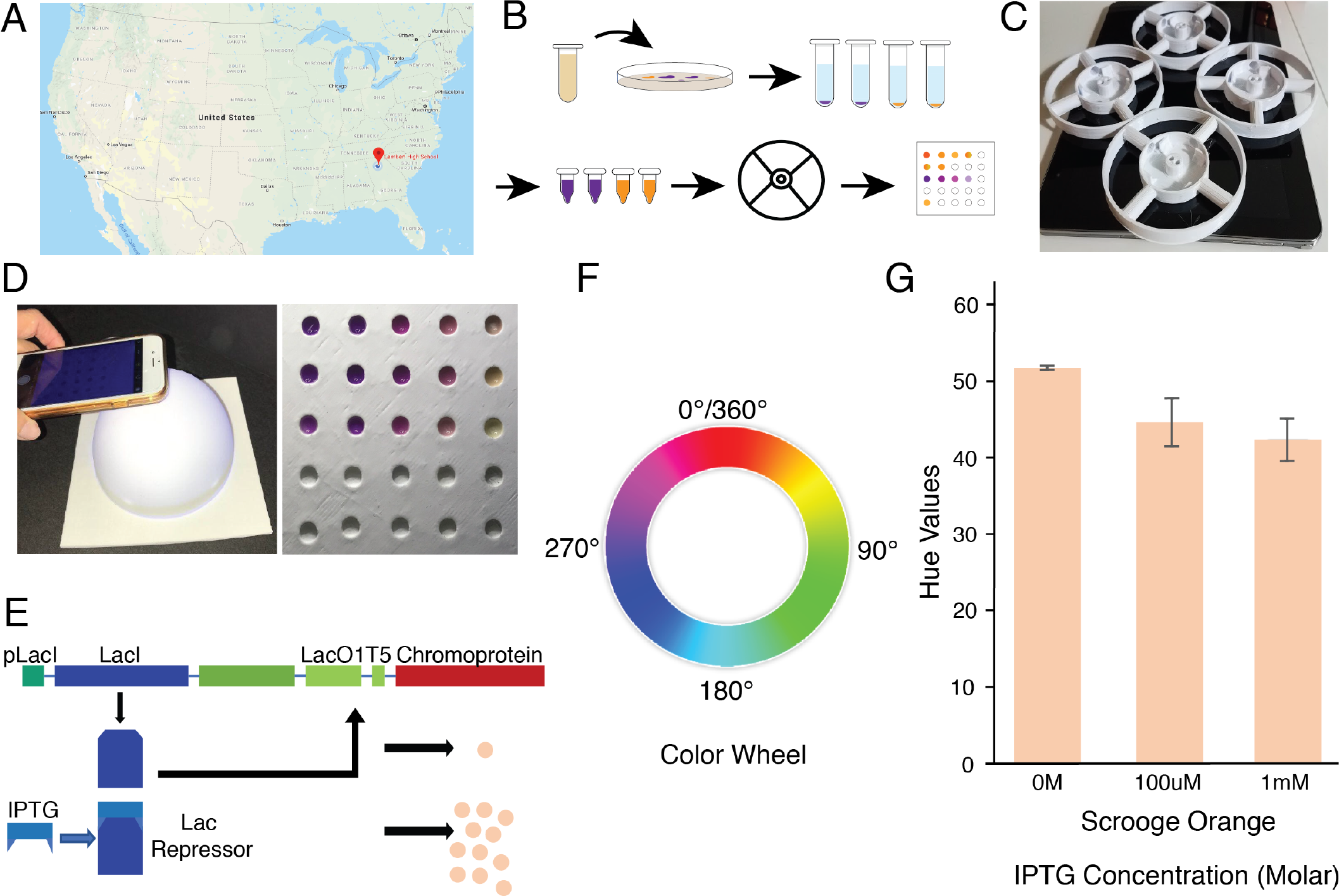
Analysis of Chromoprotein Expression in High-School. (A) Location of high school on map of United States. (B) Workflow schematic for color analysis, beginning with transformation of the plasmid and plating, growth of liquid cultures, isolation of small portion of culture, centrifugation with 3D-Fuge, and sample illumination chamber and color picker for measurements. (C) Images of 3D-Fuges utilized in experiments. (D) Image of sample illumination chamber and phone used for capturing the image (Left). Image of view of chromoprotein-expressing bacterial pellets in sample illumination chamber. (Right). This image does not represent the pellets utilized for measurements indicated in Fig 3G. (E) Diagram for regulation of chromoprotein expression in plasmid construct. (F) Color wheel utilized for hue value measurements. As darker shades occur counter-clockwise around the color wheel, darker colors correlate with a smaller hue value. (G) Hue value measurements for Scrooge Orange chromoprotein at varying IPTG concentrations. As expected, increasing IPTG concentrations results in smaller hue values, indicating successful analysis of color expression using the 3D-Fuge and sample illumination chamber. SD, n = 3 technical replicates.

The chamber was utilized to ensure standardized white illumination of the sample and facilitate the capturing of an image through the opening at the top (Fig 3D). The corresponding images of samples were then processed through an RGB color-analysis tool to obtain the hue values, signifying the level of chromoprotein expression present. Using this measurement scale, increased chromoprotein expression would be represented by a lower value whereas decreased chromoprotein expression would be represented by a higher value. This is due to the orientation of the RGB color space where specific colors increase in darkness counter-clockwise (or in order of decreasing values) (Fig 3F). From the trials conducted, the expected decrease in hue values can be seen with increase in IPTG concentration (Fig 3G), indicating that the 3D-Fuge was able to successfully centrifuge the liquid culture samples and obtain bacterial pellets reflecting the levels of chromoprotein expression. We thereby demonstrate the applications of the 3D-Fuge in enabling chromoprotein analysis for synthetic biology education in high schools (Fig 3A), opening up new possibilities for low-cost biomarker analyses that do not require expensive fluorescence microscopes for detection.

## DISCUSSION

A major obstacle towards the advancement of field biology and science education persists through the lack of affordability, accessibility, and portability for quantification and analysis of experimental samples. This presents a significant challenge towards straightforward applications such as diagnostics in the field, as well as providing opportunities to young aspiring scientists. Although industrial equipment functions optimally in a laboratory setting, the same cannot be said for implementation under remote conditions. This issue has stimulated the rise of frugal science, a new approach towards making scientific tools more accessible and transportable. The latest of the developments using these principles is the 3D-Fuge, a 3D-Printed, low-cost hand powered centrifuge, based on the principles of the paperfuge [2]. The 3D-Fuge is capable of reaching speeds of 6,000 rpm, and functions analogous to the previously demonstrated paperfuge. The need for low-cost centrifuges is vast, due to their application in a wide spectrum of fields within science. We specifically demonstrated the application of the 3D-Fuge in assisting biomarker identification and quantification, as well as for nucleotide extractions in the field.

Developments in portable nucleotide sequencing hold great promise for fields such as human health applications, metagenomics, agriculture, and molecular taxonomy [14]. Third-generation portable sequencing instruments, such as the MinION (ONT), require high quality input DNA or RNA [16]. We therefore set out to obtain purified DNA extractions without access to conventional laboratory equipment while in a remote tropical rainforest for a real-time DNA barcoding study, which can allow for the identification of specimens via DNA amplification and sequencing [10].

For this purpose, the 3D-Fuge was designed to hold DNA spin column tubes (2 mL). Spin columns are so called because reagents are added to the top of the tube and subsequently passed through a binding matrix when spun in a centrifuge. Spin columns typically use a solid silica support to which nucleic acids bind in the presence of a chaotropic salt, allowing them to be separated from cellular proteins and polysaccharides. These salts are typically removed with an alcohol-based wash and the nucleotides are eluted off the silica column in a low-ionic-strength solution such as TE buffer or water. Binding, washing and eluting the DNA can be done rapidly in this way, with the whole process taking less than 30 minutes. The current price per commercial spin column can be approximately 1 dollar on the U.S. market and this cost can potentially be reduced by recharging used commercial spin columns or assembling homemade spin columns using filter paper as binding material [20].

Interestingly, a recent study demonstrated an equipment-free protocol using untreated cellulose-based paper to rapidly capture nucleic acids and retain them during a single washing step [22]. While we have not tested this method in our hands, a centrifuge could be avoided if the dipstick methodology allows for obtaining DNA/RNA of sufficient quality from samples. For some samples that are difficult to isolate nucleotides from and perform certain experiments (such as long-range PCR or nanopore sequencing), a more robust method involving spin-columns and additional wash steps may be necessary to produce high-quality nucleic acid extracts, for which the 3D-Fuge can be applied for centrifugation steps. We found that DNA extractions using the 3D-Fuge were of sufficient quality compared to a standard benchtop centrifuge, with extracts used for downstream long-range PCR amplification of products from around 3,500 bp (spanning the ribosomal cluster) to around 9,000 bp (spanning about half of the human mitochondrial genome) in length (Fig 1, Fig 2). A limitation to the current 3D-Fuge design is that it can only hold up to four spin column tubes at once, thus restricting the number of samples that can be processed simultaneously. Due to the low-cost nature of the 3D printed device, several units can be produced and additional users can perform hand-powered centrifugation steps.

As high schools often do not have the same resources as universities or other similar laboratories, there is a need for low-cost technologies to enable synthetic biology education in such environments. As reporter proteins are commonly employed in genetic constructs, biomarker analysis is one workflow to introduce synthetic biology into schools, and the 3D-Fuge represents the next innovation in frugal science providing students with the necessary tools to conduct experiments in low-resource settings. Using the 3D-Fuge, liquid culture samples were successfully centrifuged to concentrate bacteria into pellets for subsequent measurements. The results of centrifuging the bacterial samples demonstrated the expected trend, indicating the ease of use of the device as well as the precision performance of the 3D-Fuge. With numerous other reporter proteins to be tested, the 3D-Fuge represents another step to better enable synthetic biology education in high-schools.

## CONCLUSIONS

The field of frugal science is helping to develop new low-cost portable tools and applications, such as the ability to perform real-time diagnostics in remote environments, and enabling greater access to those with an interest in scientific devices, such as high school students. Here we introduce the 3D-Fuge, a 3D-printed device capable of centrifugation of a wide variety and volumes of solutions, such as spinning down samples for biomarker applications and performing nucleotide extractions as part of a portable molecular lab setup. Overall, we hope that the design and proof-of-principle trials presented here will stimulate others to continue research into the development of low-cost scientific devices, and that the 3D-Fuge will be valuable to a range of users including students, labs in resource limited settings, and field researchers.

## MATERIALS AND METHODS

### Chromoprotein Transformation and Analysis

The chromoprotein plasmid construct (Scrooge Orange) was purchased from the corporation ATUM Bio (ATUM, 2018) to test in the competent *E. coli* cells. The samples obtained from ATUM Bio were hydrated in 10 *μ*L of ddH20, and transformed into New England Biolabs DH5a *E. coli* using the NEB High Effciency Transformation Protocol. The resulting transformation solutions were plated onto LB agar containing the antibiotic carbenicillin. Single colonies were diluted into 40 *μ*L of water, and 1 *μ*L of the colony dilution was inoculated into three different 5 mL LB broth liquid cultures with 5 *μ*L of carbenicillin (100 *μ*g/ml). These were induced with IPTG at concentrations of 0 M, 100 *μ*M and 1 mM respectively. Liquid cultures grown for 24 hours in an 37° Celsius incubator set to shake at 170 rpm, and then removed from the incubator and allowed to sit undisturbed for 1 hour. 75 *μ*L of the settled particulate was transferred into a 0.2 mL PCR tube and centrifuged for 5 minutes using the 3D-Fuge. The supernatant was discarded and the bacterial pellets were placed into the sample illumination chamber for measurements. An image captured with a phone and an RGB color-analysis tool were utilized to determine the hue values of each bacterial pellet. These were then compared with the concentration of IPTG induction for analysis.

### 3D-Fuge Design and Materials

In Case Study 1, one male individual (age 29) utilized a 3D-Fuge capable of holding microcentrifuge tubes as well as a combination of spin columns and ow-through tubes. In Case Study 2, four identical 3D-Fuges were utilized for the centrifugation of the chromoprotein samples. All devices were printed using a custom-built 3D Printer and CAD files modified for the inclusion of tube holders to ensure PCR tubes are securely placed in the device.

The 3D-Fuge design files can be found at BhamlaLab 3D-Fuge. The design for nucleotide spin column extractions consists of a base that is 60 mm in diameter, a top that can fit over the base with two screws, and four rectangular holders that slide into grooves in the base that each hold one spin column tube in place (SI Fig1A). The devices were printed using either a Lulzbot Mini or a Lulzbot Taz with Polylite (PLA) 2.85 mm filament.

The string used in this study was the Dorisea Extreme Braid 500 lb 2.0 mm Fishing Line. String lengths were maintained at a standard 104 cm, and 3D-Fuges were coiled 50 times prior to centrifugation. We also recommend the use of safety goggles when using the 3D-Fuge as the device and components can reach high speeds.

### Nucleotide Extractions

The 3-D centrifuge was used to perform DNA extractions using the Quick-DNA Miniprep Plus kit (Zymo Research) according to the protocol. For DNA extraction comparisons in the lab, a benchtop centrifuge (Eppendorf model 5414 D) was used and each centrifugation step was performed at 13,000 rpm for 1 minute. Solid tissues collected in the field were lysed and added to a microcentrifuge tube with 95 *μ*L of water, 95 *μ*L of solid tissue buffer, and 10 *μ*L of protinase K for 1-3 hours. 400 *μ*L of Genomics Binding Buffer was added to the supernatant and transferred to a Zymo-spin Column in a collection tube. The tubes were placed into the 3D-Fuge and spun by hand for approximately one to two minutes. The flow through was discarded, then 400 *μ*L of DNA pre-wash buffer was added and centrifuged using the 3D fuge. Two rounds of washing steps were performed using gDNA Wash Buffer with 3D fuge centrifugation steps for one minute each followed by discarding of the flow through. 30 *μ*L of DNA elution buffer was added and a final centrifugation step was carried out in a clean 1.5 *μ*L tube. The final purified DNA was used for downstream molecular experiments, including polymerase chain reaction. Primers for long-range PCR were designed to amplify mitochondrial DNA (Fig 1F)[5] or ribosomal DNA [13]. Amplicons were pooled and sequenced on the MinION platform (Oxford Nanopore Technologies)[13].

## ACKNOWLEDGEMENTS

A.F.P would like to thank staff at the Jacobs Institute for Design Innovation as well as members of the Patel Lab, Rainforest Expeditions, Helen Kurkjian, Bjorn Hartmann and the Center for Interdisciplinary Biological Inspiration in Education and Research (CiBER) at the University of California Berkeley for feedback and helpful suggestions on the project, which was funded in part by the Jacobs Institute Innovation Catalyst Award. We would also like to thank E. Kim and C. Lee for assisting with centrifugation of chromoprotein samples, and the rest of the 2018 Lambert iGEM Team for their support. M.S.B acknowledges funding support through NSF (award no. 181733) and Mindlin Foundation.

## AUTHOR CONTRIBUTIONS

MSB, AFP, and GB conceptualized and designed the research. GB and JS performed experiments for the Case Study 2 (chromoprotein analysis) and AFP performed experiments and analysis for Case Study 1 (nucleotide extractions). SS and GB captured and analyzed the high-speed footage of the 3D-Fuge. All authors analyzed the data and wrote the manuscript.

