## Supplementary Information for "A 3D-printed hand-powered centrifuge for molecular biology"

(Dated: January 13, 2019)

### CONTENTS

I. Supplementary Movies 1
A. S1: 3D-Fuge: a 3D-Printed Hand-Powered Centrifuge 1

II. Supplementary Figures 1

### I. SUPPLEMENTARY MOVIES

#### A. S1: 3D-Fuge: a 3D-Printed Hand-Powered Centrifuge

Short video demonstrating the need and applications of the 3D-Fuge both in field conditions and within high schools. The video can be viewed on Youtube at: The 3D-Fuge.

### II. SUPPLEMENTARY FIGURES

* authors contributed equally

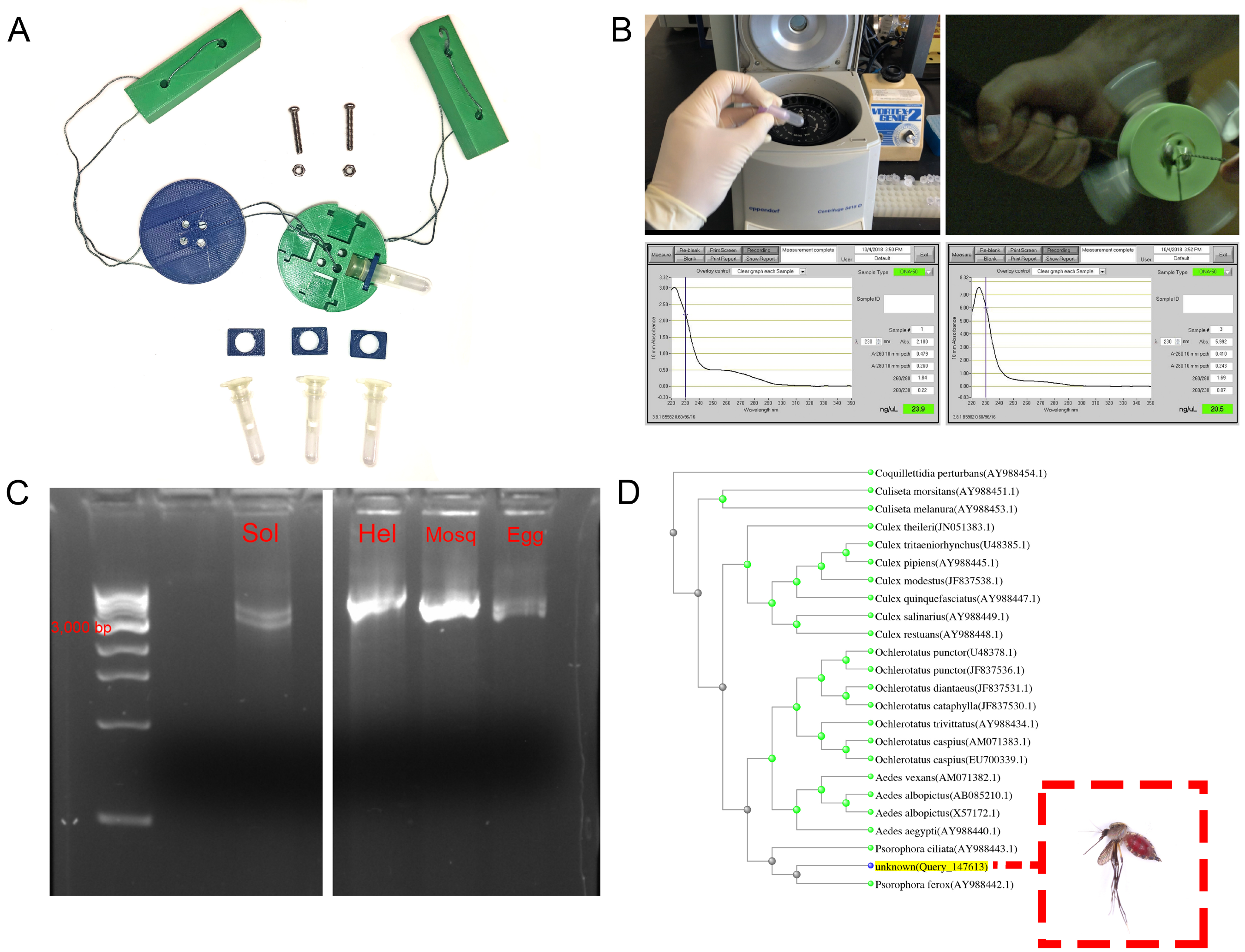


FIG. S1. **Nucletide Extractions with 3D-Fuge.**(A) Components and 3D printed parts of the 3D-Fuge. (B) Comparison of human cheek swab DNA extractions using a conventional laboratory bench top centrifuge (left) and the 3D-Fuge (right) with their respective Nanodrop DNA quantifications. Long-range mitochondrial PCR products using these extracts can be found in Figure 1E. (C) Gel electrophoresis of samples that were extracted in the field using the 3D-Fuge and subsequently PCR amplified with ribosomal DNA primers (left to right: Solanaceae, Heliconius butterfly, Mosquito, and Butterfly eggs). (D) NCBI distance of tree results from a consensus sequence generated in the field from the bloodfed mosquito sample.
